# Population-based neuroimaging reveals traces of childbirth in the maternal brain

**DOI:** 10.1101/650952

**Authors:** Ann-Marie G. de Lange, Tobias kaufmann, Dennis van der Meer, Luigi Maglanoc, Dag Alnæs, Torgeir Moberget, Gwenaëlle Douaud, Ole A. Andreassen, Lars T. Westlye

## Abstract

Pregnancy and childbirth involve maternal brain adaptations that promote attachment to and protection of the newborn. Using brain imaging and machine learning, we provide evidence for a positive relationship between number of childbirths and a ‘younger-looking’ brain in 12,021 women, which could not be explained by common genetic variation. The findings demonstrate that parity can be linked to brain health later in life.

---

During pregnancy and postpartum, fundamental biological processes are instigated to support maternal adaptation and ensure protection of the offspring [1]. In rodents, brain adaptations across pregnancy and postpartum include altered neurogenesis in the dentate gyrus [2], and changes in volume, dendritic morphology, and cell proliferation in the hippocampus [1, 3]. In humans, reduction in total brain volume has been observed during pregnancy, with reversion occurring within six months of parturition [4]. Regional changes in brain structure are evident during the postpartum period, with effects depending on region and time since delivery [5–8]. While some maternal brain changes revert postpartum, others extend well beyond this phase [1, 7–9] and may influence the course of neurobiological aging later in life. Some regional grey matter changes have been found to endure for at least 2 years post-pregnancy in humans [1], and aged parous rats have increased hippocampal long-term potentiation and show fewer signs of brain aging [1, 10]. In addition to the direct and indirect bodily and environmental adaptations in response to pregnancy and child-rearing, such long-lasting effects on brain health in humans could also reflect genetic pleiotropy, as reproductive behaviors are complex, heritable traits with a polygenic architecture that partly overlaps with a range of other traits that influence brain-health trajectories [11].

Based on the evidence of long-lasting effects of parity on the maternal brain, we investigated structural brain characteristics in 12,021 women from the UK Biobank, hypothesizing that women who had given (live) birth (n = 9568) would show less evidence of brain aging compared to their nulliparous peers (n = 2453). We used machine learning and brain age prediction to test I) if a classifier could identify women as parous or nulliparous based on morphometric brain characteristics, and II) whether brain age gap (estimated brain age – chronological age) differed between parous and nulliparous women. Mean age (± SD) was 54.72 (7.29) years for the full sample; 55.23 (7.22) years for parous and 52.79 (7.23) years for nulliparous women. To investigate the impact of number of childbirths, we tested for associations between number of births and the probabilistic scores from the group classification and brain age gap, respectively, in addition to comparing women who had given 1-2 births, 3-4 births, and 5-8 births to nulliparous women, respectively. To parse the effects of common genetic variation, we performed a genome-wide association study (GWAS) on the phenotype *number of births* in 271,312 healthy women in the UK Biobank (excluding our MRI subsample). We then computed polygenic score for each European individual in our MRI subsample (N = 10,289, Online Methods), and tested for associations between polygenic scores and the probability score from the group classification and brain age gap, respectively. To estimate genetic overlap between number of births and a range of other complex traits, we used linkage disequilibrium score regression based on previously published GWAS results [12–17].

Figure 1 and Table 1 show the results from the group classification and the brain age prediction. The probability of being classified as parous was positively related to number of births (*r* = 0.05, *p* = 2.30 × 10^−4^, CI = [−0.02, −0.08]), indicating a higher probability for multiparous women to be labeled correctly. In the brain age analysis, the correlation between predicted and chronological age was *r* = 0.61, *p* =< .0001, CI = [0.6, 0.62], and RMSE = 5.78 (SD = 0.10), *p* =< .0001. To account for age-related bias in the predicted age [18, 19], we employed a quadratic regression to the data (Equation 1, Online Methods). Bias-corrected brain age gap correlated negatively with number of births (*r* = 0.07, *p* = 5.00 × 10^−16^, CI = [−0.09, −0.06]), indicating a ‘younger looking’ brain in multiparous women. The correlation remained significant when including only parous women (*r* = 0.03, *p* = 3.14 × 10^−3^, CI = [−0.05, −0.01]). To assess the reproducibility of these effects, the brain age analysis was rerun using predicted brain age estimates based on an independent approach and training set [20–22]. In brief, the results were consistent with the main findings (see Online Methods for full description). To investigate relevant confound variables, we performed additional analyses testing the associations between brain age gap and number of childbirths when accounting for ethnic background, education, body mass index, and age at first birth. None of these variables fully explained the differences in brain age gap between parous and nulliparous women. The results are provided in Supplementary Tables 2–5.

**Figure 1:**
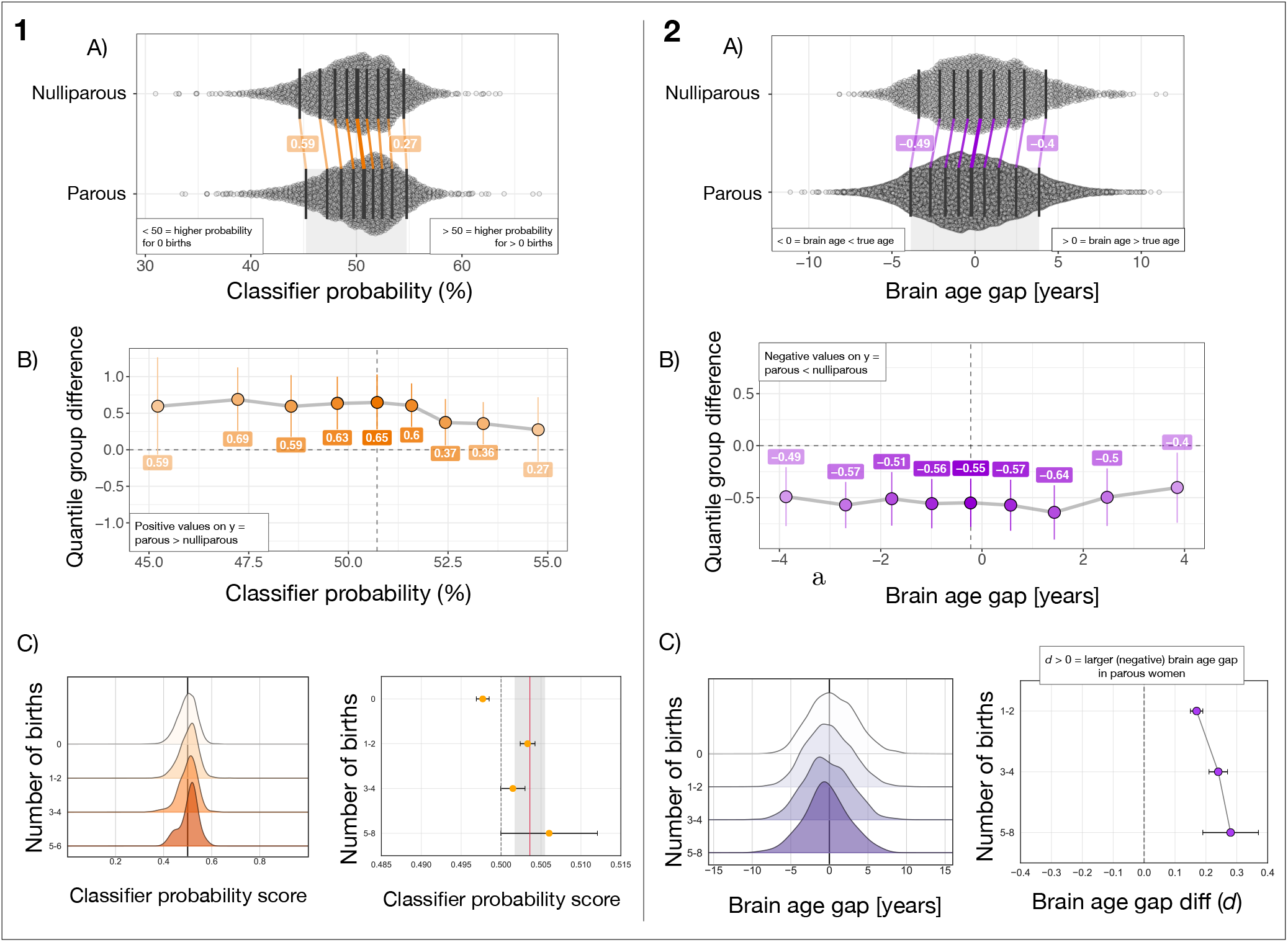
**Panel 1** A) The distributions of classifier probability scores in nulliparous and parous women. The x-axis refers to the estimated percentage probability of having given birth. The dark vertical lines mark the deciles of the distributions. The matching deciles in the two groups are joined by colored lines, showing a uniform, positive shift in the group of parous women. B) The portion of the x-axis in plot A marked by the grey shaded area at the bottom of the plot. The y-axis shows the group differences between deciles (parous group minus nulliparous group), while the x-axis shows the deciles of the parous group. C) Left plot: The distribution of individual-level classifier probability scores in subgroups of women based on number of childbirths. The plot shows a positive shift in the distribution with a larger number of births. The plot is displayed with balanced group samples (n nulliparous women = 2453, 1-2 births = 1773, 3-4 births = 645, and 5-6 births = 35, see Online Methods for details). Darker color indicates a larger number of births. Right plot: Mean classifier probability for each of the subgroups. The red vertical line shows the mean classifier probability in the groups of parous women. The lighter grey colored area illustrates the standard deviation. The error bars represent the standard error on the means. The dashed line indicates 0.5 on the x-axis. **Panel 2** A) The distributions of bias corrected, estimated brain age gap in nulliparous and parous women. Negative values indicate a predicted brain age that is lower than chronological age, i.e. a ‘younger-looking’ brain. The plot shows a uniform, negative shift in the group of parous women. B) The y-axis shows the group differences between deciles (parous group minus nulliparous group), while the x-axis shows the deciles of the parous group. C). Left plot: The distribution of estimated brain age gap in subgroups of women based on number of childbirths. The plot shows a negative shift in the distribution with a larger number of births. Number of subjects: nulliparous women = 2453, 1-2 births = 6945, 3-4 births = 2497, and 5-8 births = 126. Darker color indicates a larger number of births. Right plot: Difference in brain age gap between each of the subgroups and nulliparous women as measured by Cohen’s d. The error bars represent the standard deviation of the effect size [23]. Higher values on the x-axis indicate a larger effect size. The dashed line indicates 0 on the x-axis.

**Table 1:** Results from the group classification and brain age prediction, including correlation analyses, logistic regression, and differences in brain age gap between the subgroups of parous women compared to nulliparous women, respectively. SE = standard error. Number of women with > 1 birth = 9568, nulliparous women = 2453.

| Classifier probability score and number of childbirths |  |  |  |  |  |  |
| --- | --- | --- | --- | --- | --- | --- |
| Pearson's $r$ | | | Logistic regression | | | |
| $r$ | $p$ | 95% CI | $\beta$ | SE | $z$ | $p$ |
| 0.05 | $2.30 \times 10^{-4}$ | 0.02, 0.08 | 2.19 | 0.73 | 3.01 | $2.69 \times 10^{-3}$ |
| Brain age gap and number of childbirths |  |  |  |  |  |  |
| $r$ | $p$ | 95% CI | $\beta$ | SE | $z$ | $p$ |
| -0.07 | $5.00 \times 10^{-16}$ | -0.09, -0.06 | -0.06 | 0.01 | -8.22 | $2.07 \times 10^{-16}$ |
| Group differences in brain age gap |  |  |  |  |  |  |
| Group | Mean diff (SD) | | $t$ | $p$ | Cohen's $d$ | |
| > 1 birth | 0.56 (3.76) | | 8.27 | $1.54 \times 10^{-16}$ | 0.15 | |
| 1-2 births (n = 6945) | 0.50 (2.98) | | 7.11 | $1.29 \times 10^{-12}$ | 0.17 | |
| 3-4 births (n = 2497) | 0.72 (2.99) | | 8.42 | $4.93 \times 10^{-17}$ | 0.24 | |
| 5-8 births (n = 126) | 0.82 (2.96) | | 3.02 | $2.53 \times 10^{-3}$ | 0.28 | |

The mean polygenic scores for number of births in each of the subgroups are shown in Figure 2. A positive correlation was found between polygenic scores and number of births (*r* = 0.09, *p* = 3.60 × 10^−21^, CI = [0.07, 0.11]). The correlation between number of births and classifier probability score persisted when partialling out polygenic scores (*r* = 0.08, *p* = 1.26 × 10^−7^, CI = [0.05, 0.11]). Polygenic scores and classifier probability scores showed a correlation of *r* = 0.04 (*p* = 0.07, CI = [−0.0, 0.08]) for parous women, and *r* = 0.00 (*p* = 0.99, CI = [−0.04, 0.04]) for nulliparous women (full sample: *r* = 0.03, *p* = 0.07, CI = [−0.0, 0.06]). Polygenic scores and brain age gap showed a correlation of *r* = 0.02 (*p* = 0.06, CI = [−0.0, 0.04]) for parous women and *r* = 0.04 (*p* = 0.10, CI = [−0.01, 0.08]) for nulliparous women (full sample: *r* = 0.03, *p* = 0.07, CI = [−0.0, 0.06]). The correlation between number of births and brain age gap also persisted when partialling out polygenic scores (*r* = −0.08, *p* = 1.75 × 10^−14^, CI = [−0.09, −0.06]). To estimate the genetic overlap between number of births and other traits including height, BMI, education, schizophrenia, bipolar disorder, and major depression, we used linkage disequilibrium score regression based on previously published GWAS results [12–17]. The results are provided in Supplementary Figure 1.

**Figure 2:**
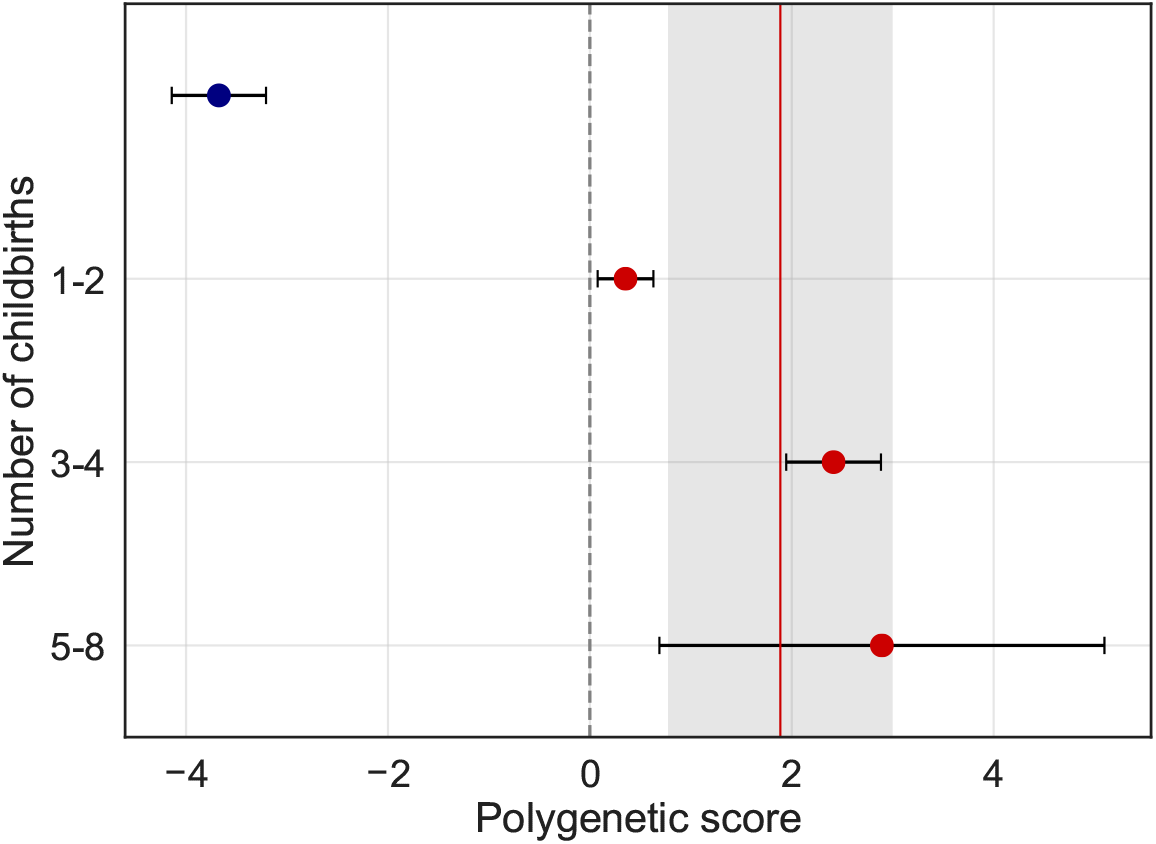
The x axis shows the mean polygenic score for number of births in each of the subgroups showed on the y-axis. Red points indicate positive scores, blue points indicate negative scores. The red vertical line and the grey shaded area show the mean polygenic score and the standard deviation in the groups of parous women. The error bars represent the standard error on the means. The dashed line indicates 0 on the x-axis.

Summarized, the results show that parity can be linked to women’s brain structure in midlife, in line with a recent analysis that tested for associations between brain age and a range of phenotypes in the UK Biobank [18]. We found no evidence that common polygenetic variation or confound variables (see Supplementary Material) could fully explain the differences in brain age gap between parous and nulliparous women. In light of the existing literature, the findings indicate that parity involves long-lasting changes in brain structure [8, 24–28] that may entail a protective effect on brain health later in life [9, 29]. Such enduring effects may also be more prominent following multiple childbirths, as multiparous women were more likely to be classified as parous based on their brain characteristics, and also had the ‘youngest-looking’ brains in terms of brain age gap.

Endocrinological modulations play an important role in the increased brain plasticity that occur during and after pregnancy [26, 28]. Changes in sex steroid hormones are known to influence human brain structure through regulation of neuronal morphology [30–32], and hormones such as estradiol, progesterone, prolactin, oxytocin, and cortisol are known to regulate brain plasticity [26, 30]. Hormonal profiles are thus likely to contribute to maternal brain adaptations during pregnancy and postpartum, and their fluctuations may have long-term implications for brain health. Lifetime duration of endogenous exposure to estrogen, which has neuroprotective effects, has been linked to risk for Alzheimer’s disease (AD) later in life [33].

Another proposed mechanism for enduring effects is the long-lasting presence of fetal cells in the maternal body [34–36], and such fetal microchimerism provides an avenue for biological interactions between fetal and maternal cells long after delivery. In an evolutionary framework, this has been conceptualized as a mother-offspring negotiation [35], providing an intriguing link to the maternal immune system. There is strong evidence for a crucial role of immune factors in pregnancy [37], which represents a state of low-level inflammation characterized by a balance between anti-inflammatory and pro-inflammatory cytokines [1, 38]. Pregnancy is known to influence and modify inflammatory disease activity and symptomology in conditions such as multiple sclerosis, asthma, and rheumatoid arthritis [39], and the pregnancy-induced increase in concentration of regulatory T cells may have implications for inflammatory susceptibility later in life. A higher cumulative time spent pregnant in the first trimester, which is when the proliferation of regulatory T cells is highest, has been shown to protect against AD [39], of which the pathogenesis is known to involve inflammatory processes [40]. Genetic differences have also been shown to interact with age and parity to influence AD neuropathology, cognitive function, and expression of proteins related to synaptic plasticity in mice [41], indicating differential genotype-dependent effects of parity on brain health across life.

In conclusion, our results provide evidence that parity is linked to brain health in midlife, and that this association cannot be explained by common genetic variability. Parity may thus involve neural changes that extend beyond the postpartum period and confer a protective effect on the aging brain.

## Acknowledgements

This research has been conducted using the UK Biobank Resource under Application Number 27412, and was supported by the Research Council of Norway (273345, 249795, 276082), the South-East Norway Regional Health Authority (2015073), and the European Research Council (ERC StG 802998).

## Online Methods

### Sample

The sample was drawn from the UK Biobank (www.ukbiobank.ac.uk), and included 12,021 women. Sample demographics are provided in Table 2 and 3.

**Table 2:** Demographics for each group. M ± SD = mean ± standard deviation. Ethnic background: W = white, B = black, M = mixed, A = Asian, C = Chinese, O = other. Educational qualification: U = university/college degree, A = A levels or equivalent, O = O levels/GCSE or equivalent, C = CSE or equivalent, N = NVQ/HNS/HNS or equivalent, P = professional qualification, e.g. nursing/teaching. For the categories, see http://biobank.ctsu.ox.ac.uk/crystal/coding.cgi?id=100305 and http://biobank.ctsu.ox.ac.uk/crystal/coding.cgi?id=1001

| Births | N | Age (M ± SD) | Ethnic background % | Educational qualification % |
| --- | --- | --- | --- | --- |
| 0 | 2453 | 52.79 (7.23) | W97.02\|B0.78\|M0.73\|A0.65\|C0.37\|O0.37 | U51.90\|A13.58\|O20.42\|C3.67\|N2.41\|P4.81 |
| >1 | 9568 | 55.23 (7.22) | W97.55\|B0.52\|M0.46\|A0.68\|C0.36\|O0.40 | U40.52\|A14.14\|O23.06\|C4.54\|N3.58\|P6.26 |
| 1-2 | 6945 | 54.90 (7.17) | W97.65\|B0.53\|M0.46\|A0.59\|C0.42\|O0.30 | U40.58\|A14.27\|O23.31\|C4.84\|N3.50\|P5.86 |
| 3-4 | 2497 | 56.10 (7.30) | W97.28\|B0.52\|M0.48\|A0.92\|C0.20\|O0.60 | U40.45\|A13.86\|O22.39\|C3.80\|N3.80\|P7.17 |
| 5-8 | 126 | 56.33 (6.96) | W97.62\|B0.00\|M0.00\|A0.79\|C0.00\|O1.50 | U38.89\|A12.70\|O22.22\|C2.38\|N3.97\|P10.32 |

**Table 3:** Age at first birth and years since first birth (M ± SD) for each group, including only subjects with available data.

| Births | N | Age at first birth \| Years since first birth |
| --- | --- | --- |
| >1 | 7935 | 26.09 (4.52) \| 29.43 (9.21) |
| 1-2 | 5313 | 26.81 (4.54) \| 28.42 (9.23) |
| 3-4 | 2496 | 24.72 (4.11) \| 31.38 (8.89) |
| 5-8 | 126 | 22.94 (3.70) \| 33.38 (7.24) |

### MRI processing

Raw T1-weighted MRI data for all participants were processed using a harmonized analysis pipeline, including automated surface-based morphometry and subcortical segmentation as implemented in FreeSurfer 5.3 [42]. In line with a recent large-scale implementation [43], we utilized a fine-grained cortical parcellation scheme [44] to extract cortical thickness, area, and volume for 180 regions of interest per hemisphere, in addition to the classic set of subcortical and cortical summary statistics from FreeSurfer [42]. This yielded a total set of 1118 structural brain imaging features (360/360/360/38 for cortical thickness/area/volume, as well as cerebellar/subcortical and cortical summary statistics, respectively).

To remove outliers, the Euler numbers [45] were extracted from FreeSurfer and averaged across the left and right hemispheres. The average values were then residualized with respect to age and scanning site using linear models, before subjects with average Euler numbers of SD ± 4 were identified and excluded (n = 109). In addition, subjects with SD ± 4 on the global MRI measures mean cortical or subcortical gray matter volume were excluded (n = 10 and n = 12, respectively), yielding a total of 12,021 subjects for the main analyses.

As a data quality cross check, the main analyses (binary classification and brain age prediction) were re-run using MRI data that was first residualized with respect to the average Euler numbers in addition to the other covariates. In brief, the results were consistent with the main findings (see Supplementary Table 1 for full results).

For the binary classification, we residualized all variables with respect to age, scanning site, ethnic background, education, and ICV using linear models. For the brain age prediction, we residualized all variables with respect to scanning site, ethnic background, education, and ICV using linear models.

### Principal component analysis

Principal component analyses (PCA) were run with z-transformed MRI variables *z* = (*x* − *μ*)/*σ*, where *x* is an MRI variable of mean *μ* and standard deviation *σ*). The top 100 components were used in the subsequent analyses, explaining 56.77% of the total variance for the classifier variables, and 56.78% for the brain age prediction variables, as shown in Figure 3. As a cross check, the correlations between number of births and a) classifier prediction value and b) brain age gap were re-run with 200 components, explaining 71.62% and 70.98% of the total variance, respectively. With 200 components included, the correlation between number of births and classifier prediction value was *r* = 0.04, *p* = 0.01, CI = [0.01, 0.06], while the correlation between number of births and brain age gap showed *r* = −0.07, *p* = 2.65 × 10^−14^, CI = [−0.09, −0.05]. As the results were consistent, 100 components were chosen to reduce computational time.

**Figure 3:**
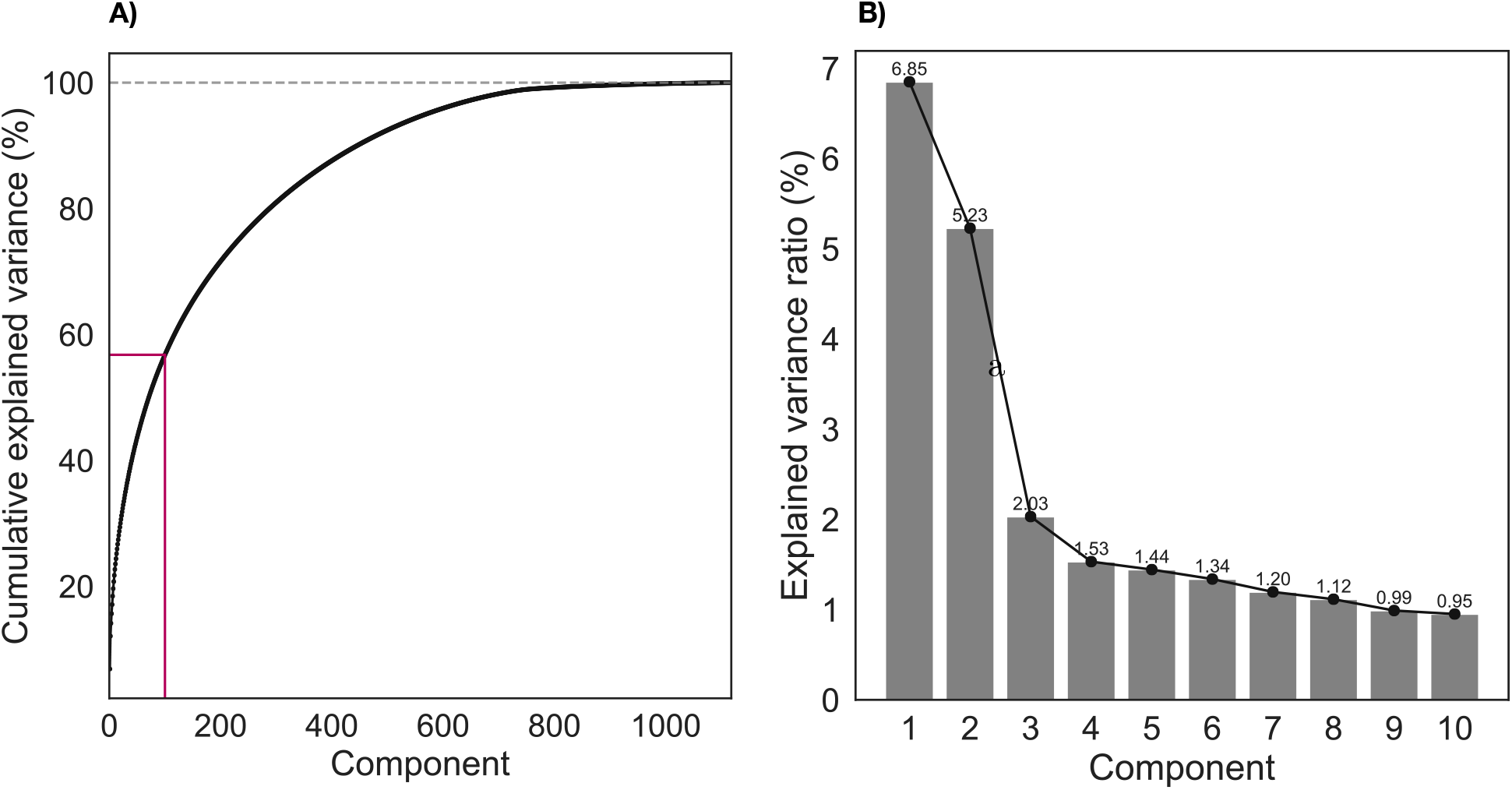
A) Cumulative explained variance for the PCA components based on 1118 z-transformed MRI variables used in the brain age analysis. B) Explained variance ratio shown for the top 10 PCA components used in the brain age analysis.

### Binary classification

Gradient boosting classification was performed using Scikit-learn (https://scikit-learn.org). Parameters were set to max depth = 1, number of estimators = 100, and learning rate = 0.1 (defaults). To account for differences in group size, under-sampling was performed using Imbalanced-learn (https://imbalanced-learn.readthedocs.io/en/stable/user_guide.html), randomly selecting samples (without replacement) in order to balance the group size. The classifier probability score was estimated based on a 10-fold cross validation, assigning a probability of being labeled as parous (having given birth) to each of the subjects. An average AUC value was generated in the cross validation, and then compared to a null distribution based on 10,000 permuted datasets. The result is shown in Figure 4.

**Figure 4:**
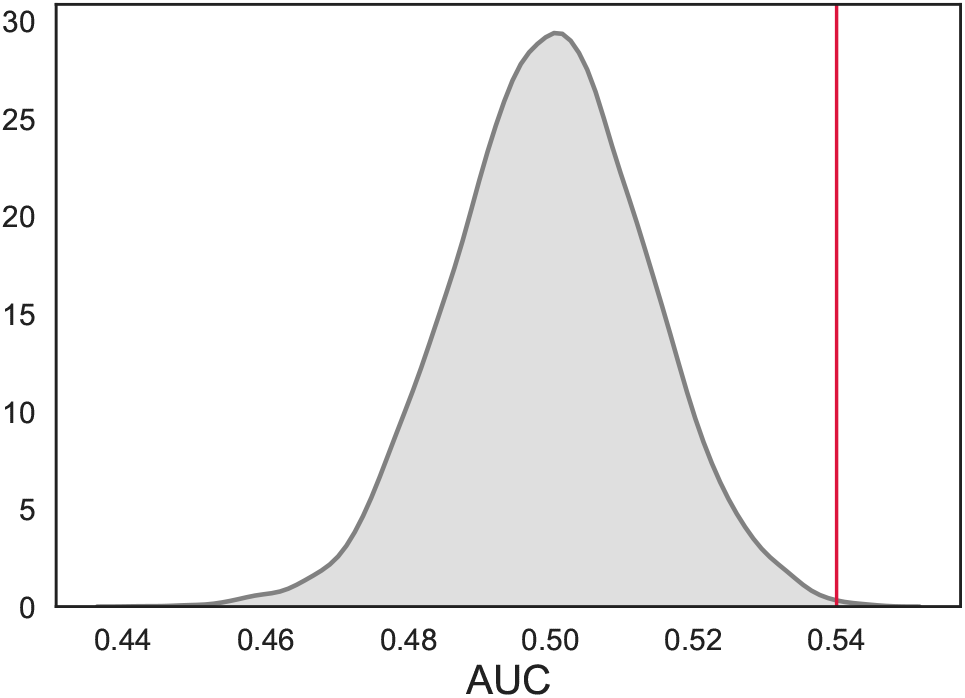
Gradient boosting classifier with PCA components based on the MRI data. AUC = 0.54 (SD ± 0.02), *p* = 6.00 × 10^−4^, based on a 10-fold cross validation (red vertical line). The null distribution calculated from 10,000 permutations is shown in gray, with an average AUC of 0.50 (SD = 0.01).

### Brain age prediction

The XGBRegressor model from XGBoost was used to run the brain age prediction analysis (https://xgboost.readthedocs.io/en/latest/python/index.html). Parameters were set to max depth = 3, number of estimators = 100, and learning rate = 0.1 (defaults). The predicted age based on the PCA components was estimated in a 10-fold cross validation with 10 repetitions per fold, assigning an estimated brain age to each individual. Brain age gap was calculated using estimated brain age - true age. The average RMSE was 5.80, as shown in Figure 5.

**Figure 5:**
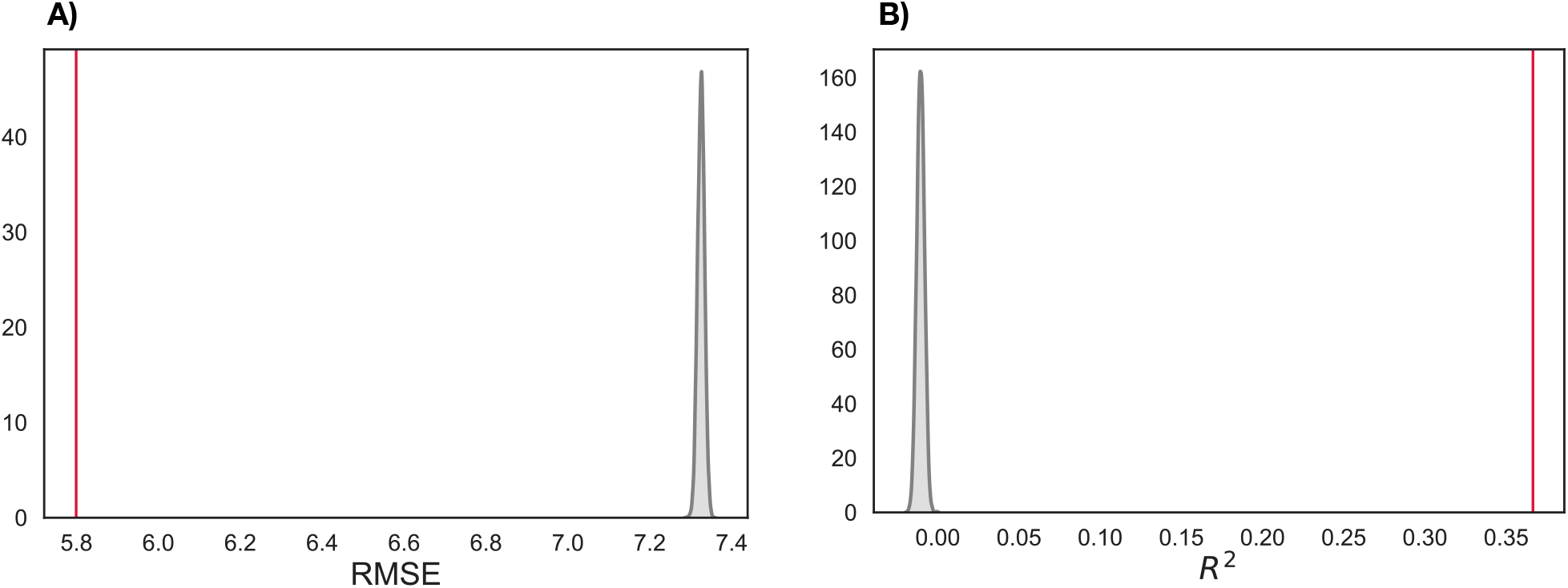
A) Average RMSE ± SD = 5.78 ± 0.10, *p* < 0.0001, based on a 10-fold cross validation with 10 repetitions per fold (red vertical line). The null distribution calculated from 10,000 permutations is shown in gray, with an average RMSE of 7.325 ± 0.009. B) Average R^2^ = 0.37 ± 0.02, *p* < 0.0001, based on a 10-fold cross validation with 10 repetitions per fold (red vertical line). The null distribution is shown in gray, with an average R^2^ of −0.011 ± 0.002.

In order to adjust for a frequently observed bias leading to generally overestimated age predictions at low age and underestimated predictions at high age [18, 19], we employed the following regression:

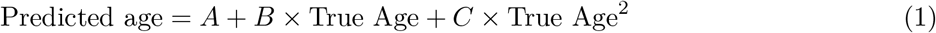

where the coefficients *A, B* and *C* parameterize the relationship between the true and predicted age. These coefficients were then used to remove the effect of the bias, in order to achieve a linear dependence with slope = 1 between the true and predicted age values, as illustrated in Figure 6.

**Figure 6:**
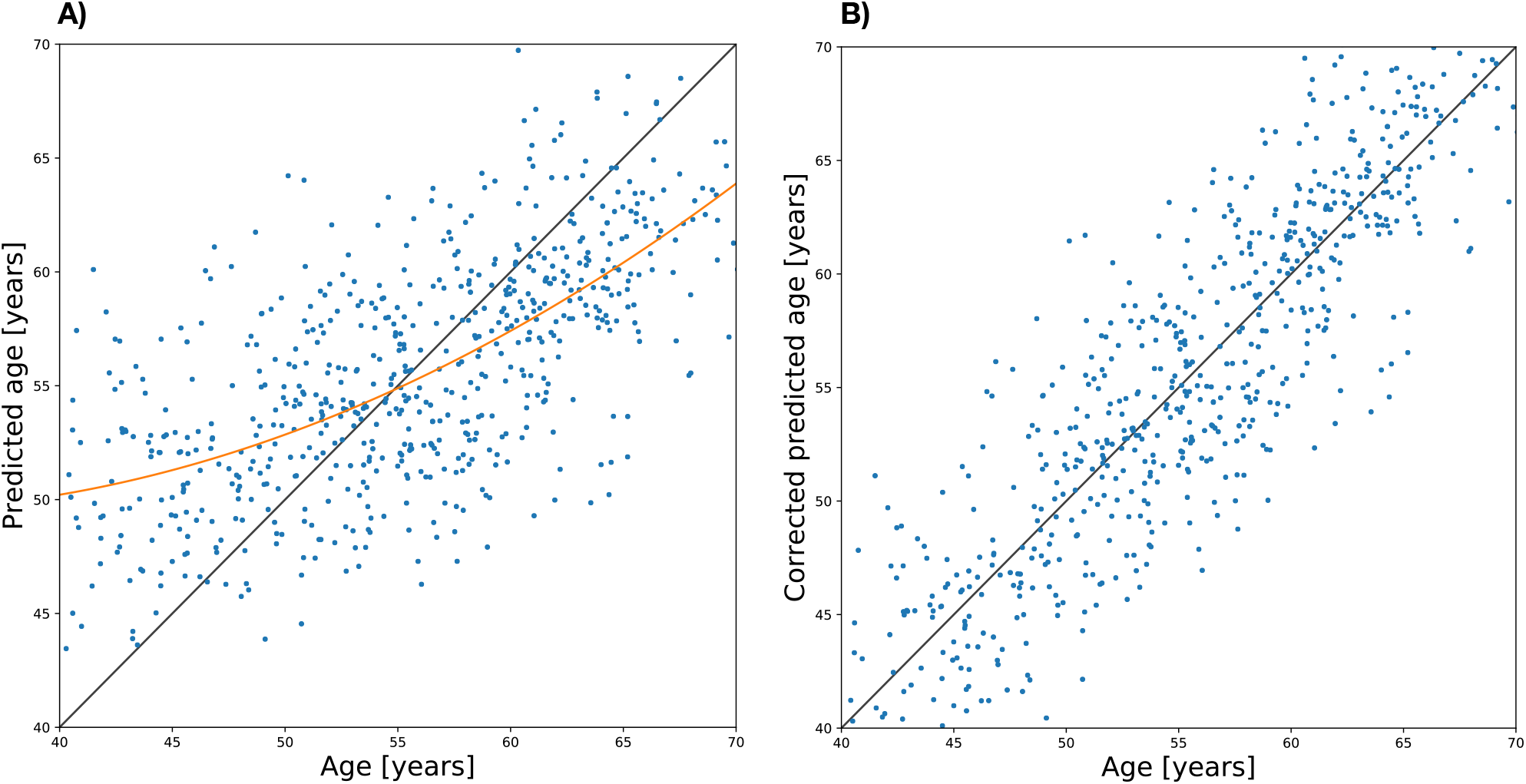
A) Machine performance is biased towards the mean age, resulting in overestimated predictions at low age and underestimated predictions at high age. B) After bias correction using Eq. 1, the predictions follow the expected dependence.

To assess the reproducibility of the effects, the brain age analysis was rerun using predicted brain age estimates based on an independent approach and training set from the brainageR software (https://github.com/james-cole/brainageR) [20–22]. The brainageR model is trained on voxel-based morphometry maps (VBM) based on T1-weighted MRI scans from 2001 healthy individuals (male/female = 1016/985, mean age ± SD = 36.95 ± 18.12, age range 18–90 years), and uses a Gaussian Processes regression with the kernlab package in R. See [21] for details.

The predicted brain age values from the brainageR dataset were corrected for age using Equation 1, and outliers with a value of < 0 and > 90 were removed (n = 15). The brain age that was estimated using brainageR and the brain age that was estimated using our current approach showed a correlation of *r* = 0.61, *p* =< 0001, CI = [0.60, 0.62]. When using the brain age values from the brainageR estimation, a negative correlation was found between number of childbirths and brain age gap (*r* = 0.04, *p* = 3.00 × 10^−6^, CI = [−0.06, −0.02]), and the group differences showed 0.72 (SD = 8.05) years for > 1 births (*t* = 4.98, *p* = 6.52 × 10^−7^, *d* = 0.09), 0.61 (SD = 6.33) years for 1-2 births (*t* = 4.10, *p* = 4.31 × 10^−5^, *d* = 0.10), 0.92 (SD = 6.40) years for 3-4 births (*t* = 5.08, *p* = 3.95 × 10^−7^, *d* = 0.14), and 2.50 years (SD = 6.48) for 5-8 births, *t* = 4.18, *p* = 3.00 × 10^−5^, *d* = 0.38), relative to nulliparous women, respectively. The results are shown in Figure 7.

**Figure 7:**
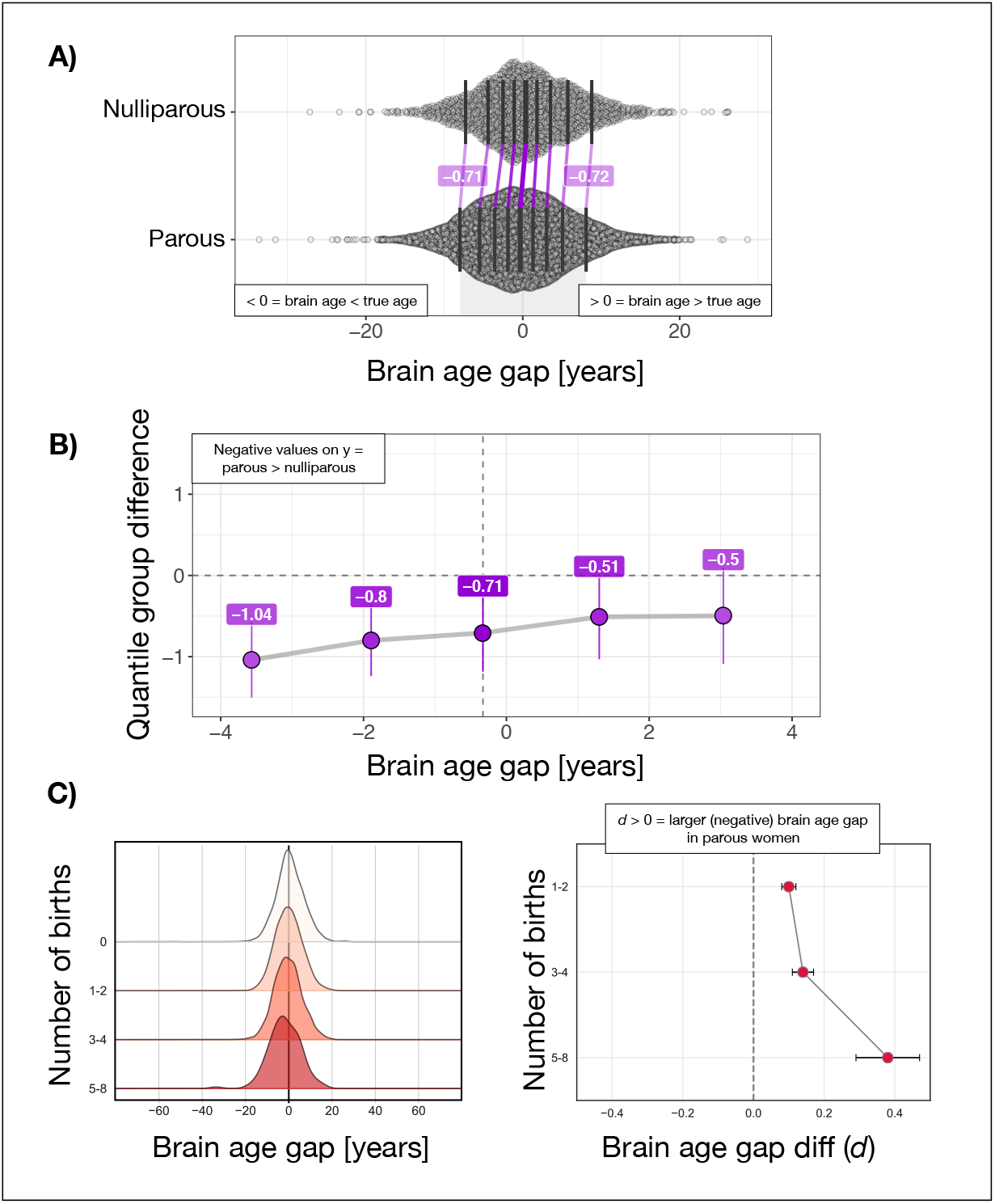
A) The distributions of bias corrected brain age gap in nulliparous and parous women. Negative values indicate a predicted brain age that is lower than chronological age, i.e. a ‘younger-looking’ brain. The plot shows a uniform, negative shift in the group of parous women. B) The y-axis shows the differences between deciles (parous group minus nulliparous group), while the x-axis shows the deciles of the parous group. C) The distribution of brain age gap in subgroups of women based on number of childbirths. The plot shows a negative shift in the distribution with a larger number of births. Number of subjects: nulliparous women = 2453, 1-2 births = 6945, 3-4 births = 2497, and 5-8 births = 126. Darker color indicates a larger number of births. C). Left plot: The distribution of estimated brain age gap in subgroups of women based on number of childbirths. The plot shows a negative shift in the distribution with a larger number of births. Darker color indicates a larger number of births. Right plot: Difference in brain age gap between each of the subgroups and nulliparous women as indexed by Cohen’s *d*. The error bars represent the standard deviation of the effect size [23]. Higher values on the x-axis indicate a larger effect size. The dashed line indicates 0 on the y-axis.

The differences between the quantiles of the groups of parous and nulliparous women were investigated using the shift function in the Robust Graphical Methods For Group Comparisons package in R [46] (https://github.com/GRousselet/rogme). In brief, the shift function shows the difference between the quantiles of two groups as a function of the quantiles of one group.

### Genome-wide association study (GWAS)

A GWAS was run on the women in the UK Biobank cohort (n = 271,312, excluding the MRI subsample), using PLINK 2.0 [47] and the UKB v3 imputed genetic data, filtering out SNPs with a minor allele frequency below 0.001 or failing the Hardy-Weinberg equilibrium test at *p* < 1.00 × 10^−9^. Non-Caucasians and individuals with a brain disorder as indicated by ICD10 were excluded from the study. We then ran a linear regression on the continuous measure number of childbirths, covarying for age and the first ten genetic principal components, as provided by UK Biobank under field 22009. PRSice v1.25 [49] was used to calculate polygenic scores for number of births across *p*-value thresholds from 0.001 to 0.5, with intervals of 0.001, using PRSice default settings. This includes the removal of the major histocompatibility complex (MHC; chromosome 6, 26-33Mb) and thinning of SNPs based on linkage disequilibrium and *p*-value. A PCA was run on the polygenic scores across all *p*-value levels [50], and the first component explaining 92.24% of the total variance was used in the subsequent analyses. The PCA component correlated *r* = 0.9 with the PGS at threshold *p* = .05. Cross-trait linkage disequilibrium score regression [48] was applied to calculate genetic correlation between number of children and height, BMI, education, major depressive disorder, schizophrenia, and bipolar disorder [12–17].

## Supplementary material

**Supplementary Figure 1:**
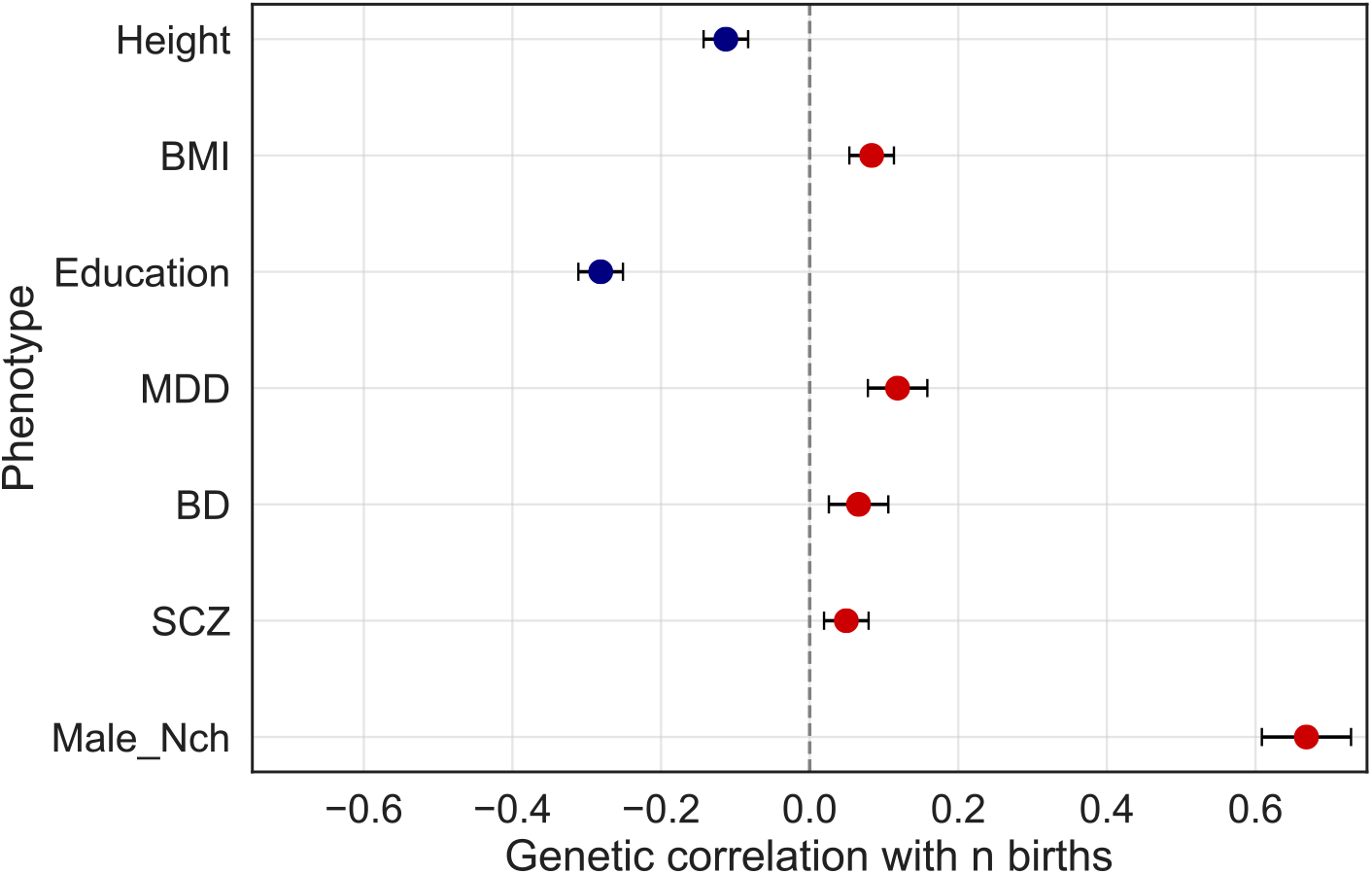
Genetic overlap between number of childbirths and other traits. Red points indicate positive correlation, blue points indicate negative correlation. The error bars indicate the standard error. BMI = Body mass index, MDD = Major depressive disorder, BD = Bipolar disorder, SCZ = Schizophrenia, Male_Nch = male number of children fathered. Linkage disequilibrium score regression revealed genetic correlations between number of childbirths and height (*r_g_* = −0.11 (SE = 0.03), *p* = 1.31 × 10^−5^), BMI (*r_g_* = 0.08 (SE = 0.03), *p* = 5.40 × 10^−3^), education (*r_g_* = −0.28 (SE = 0.03), *p* = 4.32 × 10^−28^), and major depressive disorder (*r_g_* = 0.12 (SE = 0.04), *p* = 9.15 × 10^−4^). The genetic correlation between parity and bipolar disorder (*r_g_* = 0.07 (SE = 0.04), *p* = 0.11), and schizophrenia (*r_g_* = 0.05 (SE = 0.03), *p* = 0.14) did not survive Bonferroni correction (*p* threshold = 0.007). A genetic correlation of *r_g_* = 0.67 (SE = 0.06), *p* = 9.30 × 10^−30^ was found for number of births in women and number of children fathered in men.

**Supplementary Table 1:** Correcting for Euler numbers to control for data quality. The classification and brain age prediction analyses were re-run using MRI data that were residualized with respect to the Euler numbers in addition to the other covariates using linear models. Outliers were identified and removed using the procedure described in Online Methods. The top 100 variables from a PCA were included in the regressor. The table shows the results from correlation analyses and logistic regression, and differences in brain age gap between each group of parous women compared to nulliparous women, respectively. The estimated brain age was corrected for age using Equation 1 in the Online Methods. Number of women with > 1 birth = 9568, nulliparous women = 2453.

| Classifier probability score and number of childbirths |  |  |  |  |  |  |
| --- | --- | --- | --- | --- | --- | --- |
| Pearson's $r$ | | | Logistic regression | | | |
| $r$ | $p$ | 95% CI | $\beta$ | SE | $z$ | $p$ |
| 0.06 | $8.00 \times 10^{-6}$ | 0.04, 0.09 | 2.18 | 0.70 | 3.11 | $1.86 \times 10^{-3}$ |
| Brain age gap and number of childbirths |  |  |  |  |  |  |
| $r$ | $p$ | 95% CI | $\beta$ | SE | $z$ | $p$ |
| -0.08 | $1.62 \times 10^{-16}$ | -0.09, -0.06 | -0.06 | 0.01 | -8.60 | $7.76 \times 10^{-18}$ |
| Group differences in brain age gap |  |  |  |  |  |  |
| Group | Mean diff (SD) | | $t$ | $p$ | Cohen's $d$ | |
| > 1 birth | 0.60 (3.86) | | 8.66 | $5.45 \times 10^{-18}$ | 0.16 | |
| 1-2 births (n = 6945) | 0.55 (3.06) | | 7.67 | $1.94 \times 10^{-14}$ | 0.18 | |
| 3-4 births (n = 2497) | 0.72 (3.07) | | 8.30 | $1.39 \times 10^{-16}$ | 0.24 | |
| 5-8 births (n = 126) | 0.85 (3.06) | | 3.03 | $2.49 \times 10^{-3}$ | 0.28 | |

**Supplementary Table 2:** Ethnic background. The brain age analysis was re-run on white subjects only. Outliers were identified and removed using the procedure described in Online Methods. The top 100 variables from a PCA were included in the regressor. The estimated brain age was corrected for age using Equation 1 in the Online Methods. The table shows the results from correlation analyses and logistic regression for the white subsample. Number of women with > 1 birth = 9330, nulliparous women = 2378.

| Brain age gap and number of childbirths |  |  |  |  |  |  |
| --- | --- | --- | --- | --- | --- | --- |
| $r$ | $p$ | 95% CI | $\beta$ | SE | $z$ | $p$ |
| -0.07 | $2.34 \times 10^{-14}$ | -0.09, -0.05 | -0.06 | 0.01 | -7.73 | $1.06 \times 10^{-14}$ |
| Group differences in brain age gap |  |  |  |  |  |  |
| Group | Mean diff (SD) | | $t$ | $p$ | Cohen's $d$ | |
| > 1 birth | 0.53 (3.77) | | 7.77 | $8.31 \times 10^{-15}$ | 0.14 | |
| 1-2 births (n = 6779) | 0.48 (2.97) | | 6.75 | $1.62 \times 10^{-11}$ | 0.16 | |
| 3-4 births (n = 2428) | 0.68 (3.00) | | 7.80 | $7.75 \times 10^{-15}$ | 0.22 | |
| 5-8 births (n = 123) | 0.73 (2.98) | | 2.67 | $2.71 \times 10^{-3}$ | 0.25 | |

**Supplementary Table 3:** Education. A higher level of education was related to a lower number of childbirths (*r* = −0.096, *p* = 3.32 × 10^−26^, CI = [0.08, 0.11]). The brain age analysis was re-run within groups of women with a) university or college level education, b) A levels, c) O levels or equivalent, and d) other professional qualifications (NVQ or similar). Outliers were identified and removed using the procedure described in Online Methods. The top 100 variables from a PCA were included in the regressor. The estimated brain age was corrected for age using Equation 1 in the Online Methods. The table shows the results from correlation analyses and logistic regression within each of the educational categories.

|  |  |  |  |  |  |  |
| --- | --- | --- | --- | --- | --- | --- |
| University/college. N = 3877 parous and 1273 nulliparous women |  |  |  |  |  |  |
| **Brain age gap and number of childbirths** |  |  |  |  |  |  |
| *r* | *p* | 95% CI | $\beta$ | SE | *z* | *p* |
| -0.07 | $2.00 \times 10^{-6}$ | -0.09, -0.04 | -0.07 | 0.01 | -4.28 | $1.86 \times 10^{-5}$ |
| A level. N = 1353 parous and 333 nulliparous women |  |  |  |  |  |  |
| **Brain age gap and number of childbirths** |  |  |  |  |  |  |
| *r* | *p* | 95% CI | $\beta$ | SE | *z* | *p* |
| -0.09 | $2.85 \times 10^{-4}$ | -0.14, -0.04 | -0.07 | 0.02 | -3.63 | $2.84 \times 10^{-4}$ |
| O level or equivalent. N = 2640 parous and 591 nulliparous women |  |  |  |  |  |  |
| **Brain age gap and number of childbirths** |  |  |  |  |  |  |
| *r* | *p* | 95% CI | $\beta$ | SE | *z* | *p* |
| -0.07 | $1.29 \times 10^{-4}$ | -0.10, -0.03 | -0.06 | 0.01 | -4.02 | $5.90 \times 10^{-5}$ |
| Other professional (NVQ or similar). N = 942 parous and 177 nulliparous women |  |  |  |  |  |  |
| **Brain age gap and number of childbirths** |  |  |  |  |  |  |
| *r* | *p* | 95% CI | $\beta$ | SE | *z* | *p* |
| -0.01 | 0.70 | -0.07, 0.05 | -0.03 | 0.03 | -1.01 | 0.31 |

**Supplementary Table 4:** Body mass index (BMI). The general relationship between brain age gap and number of births persisted when correcting for BMI (*r* = −0.07, *p* = 5.73 × 10^−16^, CI = [−0.09, −0.06]). To further investigate the influence of BMI, the brain age analysis was re-run within groups of women with BMI values of a) < 18.5, b) 18.5 - 25, c) 26 - 30, and d) > 30. Outliers were identified and removed using the procedure described in Online Methods. The top 100 variables from a PCA were included in the regressor. The estimated brain age was corrected for age using Equation 1 in the Online Methods. The table shows the results from correlation analyses and logistic regression within each of the BMI categories. 65 women had BMI below 18.5 (minimum BMI = 15), constituting a group that was too small to run with PCA. In this group, all the MRI variables were included.

| BMI < 18.5. N = 53 parous and 12 nulliparous women |  |  |  |  |  |  |
| --- | --- | --- | --- | --- | --- | --- |
| Brain age gap and number of childbirths |  |  |  |  |  |  |
| $r$ | $p$ | 95% CI | $\beta$ | SE | $z$ | $p$ |
| -0.04 | 0.74 | -0.28, 0.20 | 0.002 | 0.10 | 0.02 | 0.99 |
| BMI 18.5 - 25. N = 3086 parous and 790 nulliparous women |  |  |  |  |  |  |
| Brain age gap and number of childbirths |  |  |  |  |  |  |
| $r$ | $p$ | 95% CI | $\beta$ | SE | $z$ | $p$ |
| -0.06 | $6.40 \times 10^{-5}$ | -0.10, -0.03 | -0.07 | 0.01 | -5.33 | $9.96 \times 10^{-8}$ |
| BMI 26 - 30. N = 3076 parous and 842 nulliparous women |  |  |  |  |  |  |
| Brain age gap and number of childbirths |  |  |  |  |  |  |
| $r$ | $p$ | 95% CI | $\beta$ | SE | $z$ | $p$ |
| -0.08 | $7.79 \times 10^{-7}$ | -0.11, -0.05 | -0.06 | 0.01 | -4.82 | $1.43 \times 10^{-6}$ |
| BMI > 30. N = 2351 parous and 563 nulliparous women |  |  |  |  |  |  |
| Brain age gap and number of childbirths |  |  |  |  |  |  |
| $r$ | $p$ | 95% CI | $\beta$ | SE | $z$ | $p$ |
| -0.06 | $1.58 \times 10^{-3}$ | -0.09, -0.02 | -0.05 | 0.02 | -3.29 | $1.66 \times 10^{-3}$ |

**Supplementary Table 5:**
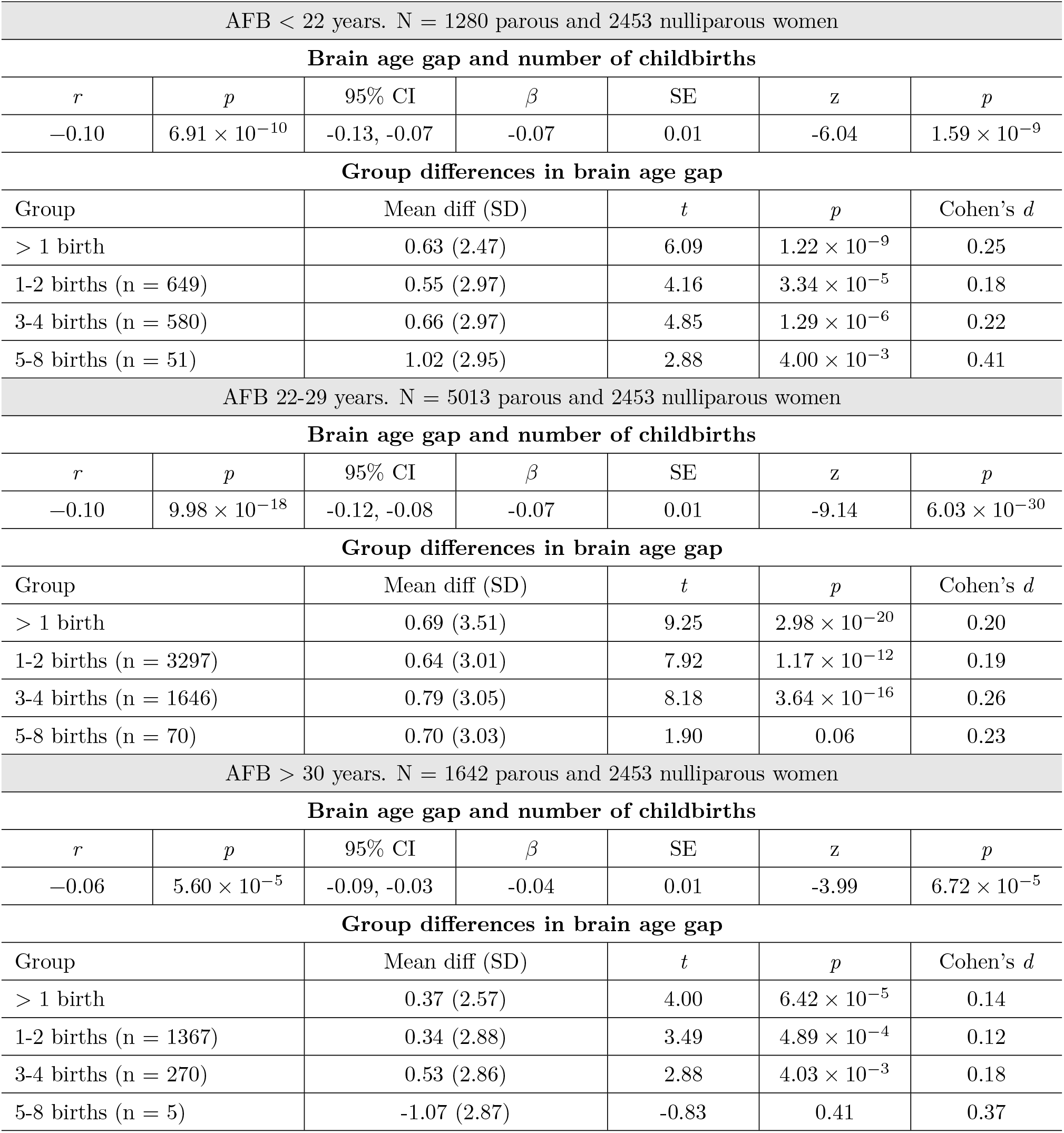
Age at first childbirth (AFB). The correlation between brain age gap and number of births was *r* = 0.02, *p* = 0.06, CI = [−0.04, −0.0]) when correcting for AFB in an analysis including only the parous women. To further investigate the influence of AFB, the brain age analysis was re-run within groups of women with AFB at a) < 22 years, b) 22 - 29 years, and c) > 30 years, as compared to nulliparous women, respectively, in a subsample including the nulliparous women and the parous women who had data on AFB (N = 8017). Outliers were identified and removed using the procedure described in Online Methods. The top 100 variables from a PCA were included in the regressor. The estimated brain age was corrected for age using Equation 1 in the Online Methods. The table shows the results from correlation analyses, logistic regression, and differences in brain age gap between each of the groups of parous women compared to nulliparous women, respectively, within each of the AFB categories.

| AFB < 22 years. N = 1280 parous and 2453 nulliparous women |  |  |  |  |  |  |
| --- | --- | --- | --- | --- | --- | --- |
| Brain age gap and number of childbirths |  |  |  |  |  |  |
| $r$ | $p$ | 95% CI | $\beta$ | SE | $z$ | $p$ |
| -0.10 | $6.91 \times 10^{-10}$ | -0.13, -0.07 | -0.07 | 0.01 | -6.04 | $1.59 \times 10^{-9}$ |
| Group differences in brain age gap |  |  |  |  |  |  |
| Group | | Mean diff (SD) | | $t$ | $p$ | Cohen's $d$ |
| > 1 birth | | 0.63 (2.47) | | 6.09 | $1.22 \times 10^{-9}$ | 0.25 |
| 1-2 births (n = 649) | | 0.55 (2.97) | | 4.16 | $3.34 \times 10^{-5}$ | 0.18 |
| 3-4 births (n = 580) | | 0.66 (2.97) | | 4.85 | $1.29 \times 10^{-6}$ | 0.22 |
| 5-8 births (n = 51) | | 1.02 (2.95) | | 2.88 | $4.00 \times 10^{-3}$ | 0.41 |
| AFB 22-29 years. N = 5013 parous and 2453 nulliparous women |  |  |  |  |  |  |
| Brain age gap and number of childbirths |  |  |  |  |  |  |
| $r$ | $p$ | 95% CI | $\beta$ | SE | $z$ | $p$ |
| -0.10 | $9.98 \times 10^{-18}$ | -0.12, -0.08 | -0.07 | 0.01 | -9.14 | $6.03 \times 10^{-30}$ |
| Group differences in brain age gap |  |  |  |  |  |  |
| Group | | Mean diff (SD) | | $t$ | $p$ | Cohen's $d$ |
| > 1 birth | | 0.69 (3.51) | | 9.25 | $2.98 \times 10^{-20}$ | 0.20 |
| 1-2 births (n = 3297) | | 0.64 (3.01) | | 7.92 | $1.17 \times 10^{-12}$ | 0.19 |
| 3-4 births (n = 1646) | | 0.79 (3.05) | | 8.18 | $3.64 \times 10^{-16}$ | 0.26 |
| 5-8 births (n = 70) |  | 0.70 (3.03) |  | 1.90 | 0.06 | 0.23 |
| AFB > 30 years. N = 1642 parous and 2453 nulliparous women |  |  |  |  |  |  |
| Brain age gap and number of childbirths |  |  |  |  |  |  |
| $r$ | $p$ | 95% CI | $\beta$ | SE | $z$ | $p$ |
| -0.06 | $5.60 \times 10^{-5}$ | -0.09, -0.03 | -0.04 | 0.01 | -3.99 | $6.72 \times 10^{-5}$ |
| Group differences in brain age gap |  |  |  |  |  |  |
| Group | | Mean diff (SD) | | $t$ | $p$ | Cohen's $d$ |
| > 1 birth | | 0.37 (2.57) | | 4.00 | $6.42 \times 10^{-5}$ | 0.14 |
| 1-2 births (n = 1367) | | 0.34 (2.88) | | 3.49 | $4.89 \times 10^{-4}$ | 0.12 |
| 3-4 births (n = 270) | | 0.53 (2.86) | | 2.88 | $4.03 \times 10^{-3}$ | 0.18 |
| 5-8 births (n = 5) |  | -1.07 (2.87) |  | -0.83 | 0.41 | 0.37 |

